# Late onset and regional heterogeneity of synaptic deficits in cortical PV interneurons of *Shank3B^−/−^* mice

**DOI:** 10.1101/2023.11.23.568500

**Authors:** Yi-Chun Shih, Lars Nelson, Michael Janeček, Rui T. Peixoto

## Abstract

Epilepsy and epileptiform patterns of cortical activity are highly prevalent in autism spectrum disorders (ASDs), but the neural substrates and pathophysiological mechanisms underlying the onset of cortical dysfunction in ASD remains elusive. Reduced cortical expression of Parvalbumin (PV) has been widely observed in ASD mouse models and human postmortem studies, suggesting a crucial role of PV interneurons (PVINs) in ASD pathogenesis. *Shank3B^−/−^* mice carrying a Δ13-16 deletion in SHANK3 exhibit cortical hyperactivity during postnatal development and reduced sensory responses in cortical GABAergic interneurons in adulthood. However, whether these phenotypes are associated with PVIN dysfunction is unknown. Using whole-cell electrophysiology and a viral-based strategy to label PVINs during postnatal development, we performed a developmental characterization of AMPAR miniature excitatory postsynaptic currents (mEPSCs) in PVINs and pyramidal (PYR) neurons of layer (L) 2/3 mPFC in *Shank3B^−/−^* mice. Surprisingly, reduced mEPSC frequency was observed in both PYR and PVIN populations, but only in adulthood. At P15, when cortical hyperactivity is already observed, both neuron types exhibited normal mEPSC amplitude and frequency, suggesting that glutamatergic connectivity deficits in these neurons emerge as compensatory mechanisms. Additionally, we found normal mEPSCs in adult PVINs of L2/3 somatosensory cortex, revealing region-specific phenotypic differences of cortical PVINs in *Shank3B^−/−^* mice. Together, these results demonstrate that loss of Shank3 alters PVIN function but suggest that PVIN glutamatergic synapses are a suboptimal therapeutic target for normalizing early cortical imbalances in SHANK3-associated disorders. More broadly, these findings underscore the complexity of interneuron dysfunction in ASDs, prompting further exploration of region and developmental stage specific phenotypes for understanding and developing effective interventions.

## Introduction

Autism spectrum disorders (ASDs) are associated with severe behavioral deficits that typically emerge during the second year of life, often following an apparently normal infancy (Lord et al. 2000; Watts 2008). The distinct developmental timing of cognitive symptom onset has long suggested that abnormal maturation of medial prefrontal cortex (mPFC) is implicated in ASD pathogenesis (Amaral, Schumann, and Nordahl 2008; Lord et al. 2000). Indeed, multiple studies have shown hypertrophy and abnormal functional connectivity of mPFC regions in autistic children (Amaral et al. 2008; Assaf et al. 2010; Gilbert et al. 2008; Goldberg et al. 2011; Sumiya et al. 2020; Turner et al. 2006; Wolff et al. 2013; Zhao et al. 2021). In addition, seizures are highly comorbid with autism, and preclinical epileptiform activity is observed in a large fraction of autistic individuals (Amiet et al. 2008; Khan et al. 2018; Lewine et al. 1999; Tuchman and Rapin 2002). However, the specific pathophysiological mechanisms underlying the onset of cortical circuit dysfunction in ASD remain elusive.

The prevalence of cortical hyperactivity in autism has led to the hypothesis that imbalances between excitatory and inhibitory (E/I) mechanisms may constitute a common pathophysiological factor in ASD (Ferguson and Gao 2018; Filice et al. 2020; Marín 2012; del Pino, Rico, and Marín 2018; Rubenstein and Merzenich 2003). In particular, deficits in the number and molecular properties of GABAergic Parvalbumin (PV) interneurons (PVINs) have been observed in several transcriptomic (Parikshak et al. 2016; Schwede et al. 2018), and postmortem histological human studies (Ariza et al. 2018; Hashemi et al. 2017). Additionally, many mouse models of autism exhibit abnormal patterns of PV expression in cortical regions (Deemyad et al. 2021; Filice et al. 2016; Gogolla et al. 2014; Negwer et al. 2020; Orefice et al. 2016, 2019; Peñagarikano et al. 2011; Pinto et al. 2014; Vogt et al. 2018; Wen et al. 2019; Zhang et al. 2020) and disrupted PVIN connectivity and function (Kalinowska et al. 2022; Negwer et al. 2020). Inhibition mediated by PVINs is a critical regulator of the activity and synchrony of local cortical circuits (Fishell and Kepecs 2020). This function relies on precise temporal recruitment during periods of heightened network activity that in turn depends on specific tuning properties of glutamatergic synapses (Akgül and McBain 2016; Jonas et al. 2004). Given that many autism risk genes are implicated in glutamatergic synapse development and function, early deficits in PVINs excitatory drive could contribute to the onset of cortical dysfunction in ASDs (Fu et al. 2022; Grove et al. 2019; Klei et al. 2012; Pinto et al. 2014; Satterstrom et al. 2019). However, identification of PVINs relies on expression of PV, that in mPFC regions only upregulated during the third postnatal week (Contractor, Ethell, and Portera-Cailliau 2021). As such, our understanding of how early PVIN function is altered by ASD risk factors, and how they might be contributing for the early pathophysiology in these conditions remains unclear.

Mutations in SH3 and Multiple Ankyrin Repeat Domains Protein 3 (SHANK3) are one of the most penetrant known causes of autism (Grove et al. 2019; Monteiro and Feng 2017; De Rubeis et al. 2018; Satterstrom et al. 2019) and are associated with seizures and profound cognitive impairments (Khan et al. 2018; Leblond et al. 2014; De Rubeis et al. 2018). SHANK3 is a complex gene that encodes a family of postsynaptic scaffolding proteins that regulate the maturation and function of glutamatergic synapses (Jiang and Ehlers 2013). Deletion of exons ι113-16 encoding the PDZ domain of SHANK3 in mice (*Shank3B^−/−^*) results in behavioral deficits and hypoconnectivity of mPFC circuits (Dhamne et al. 2017; Guo et al. 2019; Kabitzke et al. 2017; Mei et al. 2016; Pagani et al. 2019; Peça et al. 2011). In addition, *Shank3B^−/−^* mice show reduced activity of GABAergic interneurons and increased pyramidal (PYR) neuron activity in response to peripheral sensory stimulation (Chen et al. 2020; Gogolla et al. 2014; Pagano et al. 2023). Notably, lower cortical expression of PV has been one of the most replicated findings in *Shank3B^−/−^* mice (Deemyad et al. 2021; Filice et al. 2016, 2020; Gogolla et al. 2014; Orefice et al. 2019), strongly implicating PV interneuron dysfunction in SHANK3 pathophysiology. However, whether loss of Shank3 results in abnormal connectivity or function of PVINs remains to be addressed. Furthermore, cortical hyperactivity in *Shank3B^−/−^* mice is observed as early as P15, when behavioral deficits start to emerge, suggesting that abnormal maturation of PVINs might be a critical mechanism of the early pathogenesis of SHANK3 disorders (Peixoto et al. 2016, 2019). However, the postnatal maturation of PVIN synaptic and intrinsic properties is highly influenced by network activity (Alcántara, Soriano, and Ferrer 1996; Patz et al. 2004), raising questions about whether PVIN phenotypes observed in adult stages represent primary pathogenic mechanisms or secondary developmental adaptations.

To address these questions, we characterized the developmental trajectory of glutamatergic connectivity in PYR neurons and PVINs of layer 2/3 (L2/3) of mPFC in *Shank3B^−/−^* mice by whole-cell electrophysiology. To label mPFC PVINs during postnatal development we leveraged a recently developed strategy based on the S5E2 regulatory element that can specifically express transgenes in PVINs independently of the PV promoter, which, in prefrontal areas, is only upregulated during late postnatal periods (Vormstein-Schneider et al. 2020). Using this approach, we found that adult *Shank3B^−/−^*mice exhibit reduced glutamatergic innervation of both PYR and PVINs in L2/3 of mPFC, indicating that PVIN synaptic connectivity is affected by loss of Shank3. Surprisingly, these phenotypes are not observed at P15, indicating that glutamatergic synaptic deficits arise later in development, potentially as compensatory mechanisms. Moreover, contrary to what we observed in mPFC, glutamatergic innervation of PVINs in the primary somatosensory cortex of adult *Shank3B^−/−^*mice was normal, indicating that PVINs of different cortical regions exhibit specific pathophysiological phenotypes. Together, these findings demonstrate that loss of Shank3 alters glutamatergic synaptic connectivity of both PYR and PVINs L2/3 mPFC, but that these phenotypes are not observed in early postnatal development when cortical hyperactivity and behavioral deficits first arise. On a broader level, these results underscore the importance of characterizing the functional maturation of neural circuits in animal models of neurodevelopmental disorders in order to differentiate between primary mechanisms of circuit dysfunction and secondary compensatory adaptations.

## Results

### Reduced mEPSC amplitude in pyramidal neurons of L2/3 ACC of adult *Shank3B^−/−^* mice

To determine how loss of Shank3 affects glutamatergic innervation of mPFC PYR neurons we recorded AMPAR miniature excitatory post-synaptic currents (mEPSC) in acute brain slices of *Shank3B^−/−^* mice, which provide an estimate of glutamatergic synapse strength and number. We focused on the anterior cingulate cortex (ACC), given that structural and functional deficits in this region of mPFC are strongly associated with ASD (Chang et al. 2015; Langen et al. 2007, 2009, 2014; Peixoto et al. 2016, 2019; Turner et al. 2006; Willsey et al. 2013; Wolff et al. 2013), and observed in *Shank3B^−/−^* mice (Guo et al. 2019). Whole-cell recordings in L2/3 PYR neurons were performed in the presence of tetrodotoxin (TTX), a voltage-dependent sodium channel blocker, to prevent action potential propagation and evoked synaptic vesicle release, with membrane potential clamped at −70mV using a cesium based internal solution (Figure 1A). Recordings with series resistance (R_s_) > 20MΩ were discarded from analysis. PYR neurons of L2/3 ACC in adult (P60-P80) *Shank3B^−/−^* mice showed reduced mEPSC amplitude compared to *Shank3B^+/+^* (WT) littermate controls (Figure 1C, Average ± SEM PYR mEPSC amplitude in adult, WT 11.4 ± 2.4 pA; KO 10.2 ± 0.7 pA, unpaired Mann Whitney test p = 0.039, n = 26,25 neurons from n = 3,3 WT and Shank3B^−/−^ mice) whereas mEPSC frequency was similar between the two groups (Figure 1D, PYR mEPSC frequency, WT 1.7 ± 0.7 Hz; KO 2.3 ± 1.3 Hz, unpaired Mann Whitney test p = 0.224). These results indicate that in adult stages the number of glutamatergic synaptic inputs onto pyramidal neurons is not affected, but loss of Shank3 reduces synaptic AMPAR-dependent currents. Empirical cumulative distribution function (eCDF) of amplitude and frequency of randomly sampled individual mEPSCs (see methods) exhibited similar trends as cell averages but revealed a significant increase in mEPSC frequency (Figures 1I-J, Kolmogorov-Smirnov test for mEPSC amplitude p < 0.001; mEPSC inter-event interval (IEI) p = 0.007, n = 100 mEPSCs per n= 26,25 neurons from n = 3,3 WT and Shank3B^−/−^ mice). Notably, adult Shank3B^−/−^ PYR neurons also exhibited increased membrane resistance (Figure 1E, input resistance of adult PYR, WT 144.0 ± 31.9 MΩ; KO 171.8 ± 51.3 MΩ, unpaired Mann Whitney test p = 0.041). Conversely, no difference in membrane capacitance was detected in adult Shank3B^−/−^ PYR (Figure 1F, membrane capacitance of adult PYR, WT 51.0 ± 10.2 pF; KO 46.8 ± 9.8 pF, unpaired Mann Whitney test p = 0.296) as well as in mEPSC rise and decay times (Figure 1G, mEPSC rise time of adult PYR, WT 1.7 ± 0.2 ms; KO 1.7 ± 0.2 ms, unpaired Mann Whitney test p = 0.843. Figure 1H, mEPSC decay tau of adult PYR, WT 7.2 ± 0.8 ms; KO 6.9 ± 0.8 ms, unpaired Mann-Whitney test p = 0.279), suggesting that neuron cell size and AMPAR kinetics are not affected by loss of Shank3. Together, these results indicate that loss of Shank3 results in weakened glutamatergic synapses of PYR neurons in adult L2/3 ACC by decreasing either number of AMPAR or neurotransmitter content per quanta rather than changing number of functional synaptic inputs. The increased membrane resistance also suggests increased excitability of adult L2/3 PYR neurons, similar to what has been reported in other neuronal cell types in adult Shank3B^−/−^ mice (Guo et al. 2019; Wang et al. 2017; Zhu et al. 2018).

**Fig 1.**
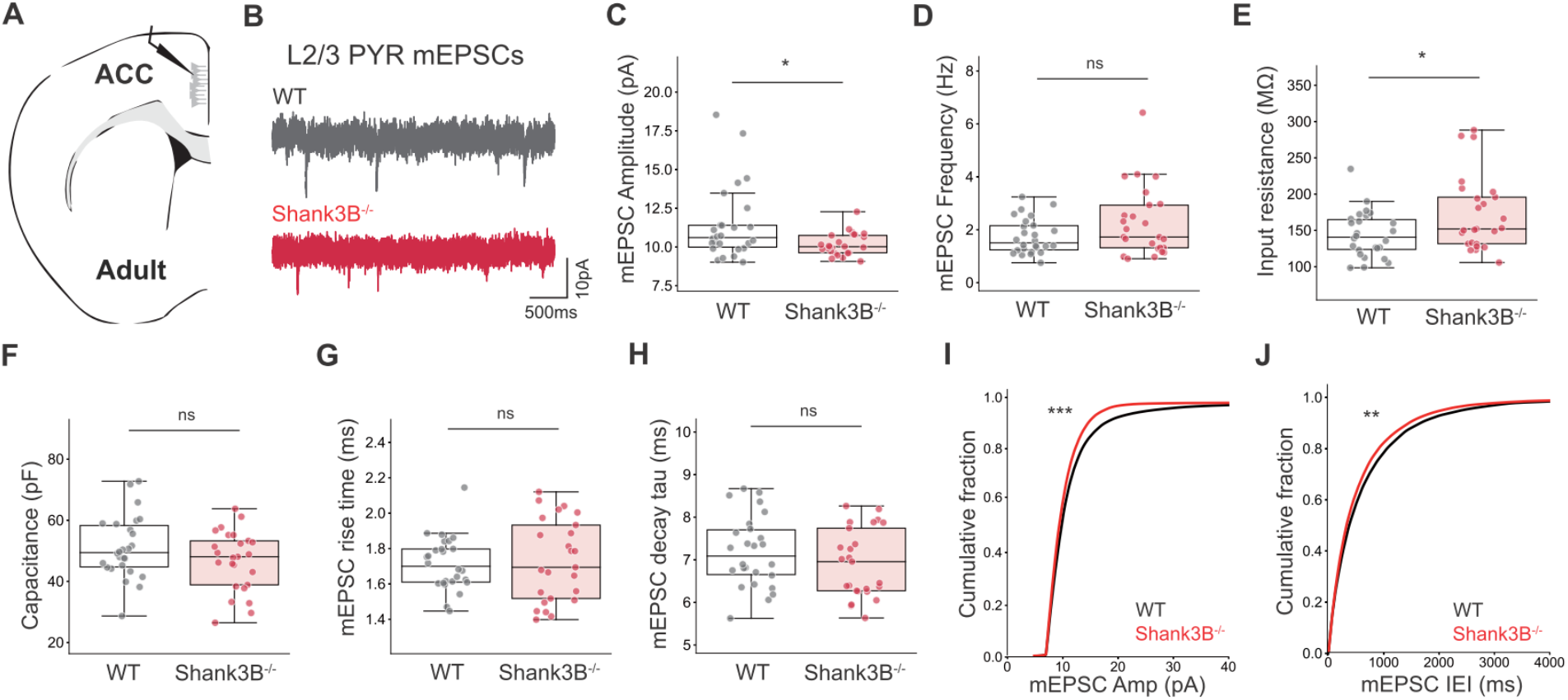
Abnormal mEPSC amplitude of PYR neurons in L2/3 ACC of adult Shank3B^−/−^ mice. (**A**) Schematic representing coronal brain section and whole-cell recordings in L2/3 PYR of adult ACC. (**B**) Representative AMPAR mEPSCs in WT and Shank3B^−/−^ mice in adult. (**C**) Average ± SD of mEPSC amplitude and (**D**) frequency I input resistant **(F)** membrane capacitance and **(G-H)** mEPSC rise and decay times of WT (gray) and Shank3B^−/−^ (red) neurons. (**I-J**) Cumulative distribution of mEPSC amplitude and mEPSC inter-event interval. n = 26,25 neurons from n = 3,3 WT and Shank3B^−/−^ mice. Box plots show median values (middle vertical bar) and ^2^5th (bottom line of the box) and ^7^5th percentiles (top line of the box) with whiskers indicating the range. eCDFs represent the average of distributions of 100 randomly sampled mEPSCs from each recorded neuron. Average mEPSC box plot *P < 0.05 with Mann–Whitney U test. mEPSC eCDFs **P < 0.01 and ***P< 0.001 with Kolmogorov-Smirnov test.

### Reduced mEPSC frequency and amplitude in PVINs of L2/3 ACC of adult *Shank3B^−/−^* mice

Adult *Shank3B^−/−^* mice exhibit reduced activity of cortical interneurons in response to sensory stimulation (Chen et al. 2020) and abnormal patterns of PV expression throughout cortex (Deemyad et al. 2021; Filice et al. 2016, 2020; Gogolla et al. 2014; Orefice et al. 2019). To determine whether loss of Shank3 results in altered glutamatergic innervation of PVINs during adult stages, we recorded mEPSCs in PVINs of L2/3 ACC of adult WT and *Shank3B^−/−^* mice previously injected (2 weeks) with AAV9-S5E2-dTom (Figure 2A). This strategy results in enriched labeling of PVINs (Vormstein-Schneider et al. 2020) that we confirmed by immunohistochemistry (IHC) against PV in ACC at P60 (Figure 2, Average ± SEM Fraction of S5E2-tdTom^+^ neurons with PV^+^ signal - 94.3 ± 4.1%, n = 125 cells from 3 mice).

**Fig 2.**
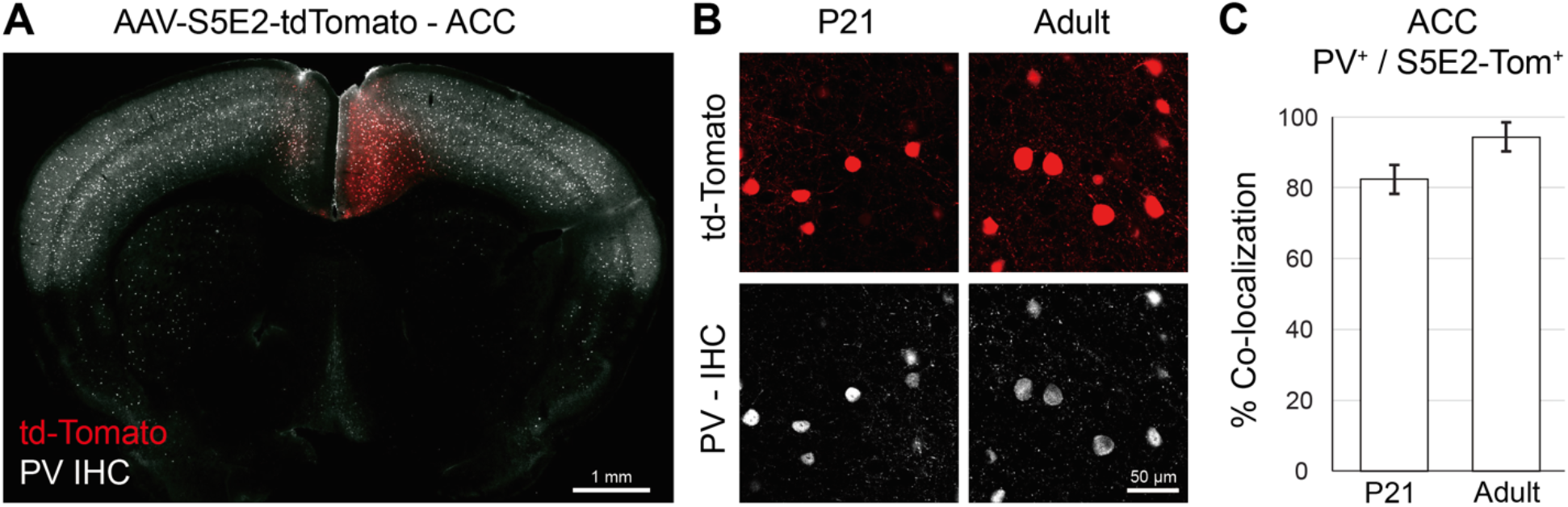
Specific labeling of PVINs with AAV-S5E2-tdTom in ACC of P21 and adult mice. (**A**) Representative image of PV IHC in ACC of P21 WT mouse injected with AAV9-S5E2-tdTom at P7 (**B**) Magnified images of PV IHC in ACC of P21 (left) and adult (right) mice after 2 weeks of injection (**C**) Percentage of tdTom-positive cells with PV co-labeling - adult ACC. n = 132,125 cells from n = 2,3 P21 and adult mice.

Voltage-clamp recordings of mEPSCs in tdTom^+^ neurons (Figure 3A and 3B) revealed that global loss of Shank3 causes a reduction of mEPSC amplitude and frequency in ACC PVINs of adult *Shank3B^−/−^* mice (Figure 3C and 3D, Average ± SD mEPSC amplitude WT 14.6 ± 2.6 pA; KO 13.1 ± 2.1 pA, unpaired Mann Whitney test p = 0.025; mEPSC frequency WT 6.3 ± 2.9 Hz; KO 5.0 ± 2.6 Hz, unpaired Mann Whitney test p = 0.019). The eCDF of individual mEPSCs showed similar reduction of amplitude and frequency in Shank3B^−/−^ PVINs compared to WT controls (Figure 3I and 3J, Kolmogorov-Smirnov test p < 0.001). Loss of Shank3 did not affect AMPAR mEPSC rise and decay kinetics, membrane capacitance or membrane resistance of adult PVINs (Figure 3E - 3H, PVIN input resistance, WT 244.6 ± 124.7 MΩ; KO 247.3 ± 77.5 MΩ, unpaired Mann Whitney test p = 0.318; PVIN membrane capacitance, WT 27.1 ± 6.5 pF; KO 29.4 ± 5.2 pF, unpaired Mann Whitney test p = 0.179; PVIN mEPSC rise time, WT 1.4 ± 0.1 ms; KO 1.5 ± 0.1 ms, unpaired Mann Whitney test p = 0.383; PVIN mEPSC decay tau, WT 2.5 ± 0.6 ms; KO 2.4 ± 0.4 ms, unpaired Mann Whitney test p = 0.353). Thus, loss of Shank3 reduces the number and strength of AMPAR excitatory input onto L2/3 PVINs and PYR neurons in adult ACC.

**Fig 3.**
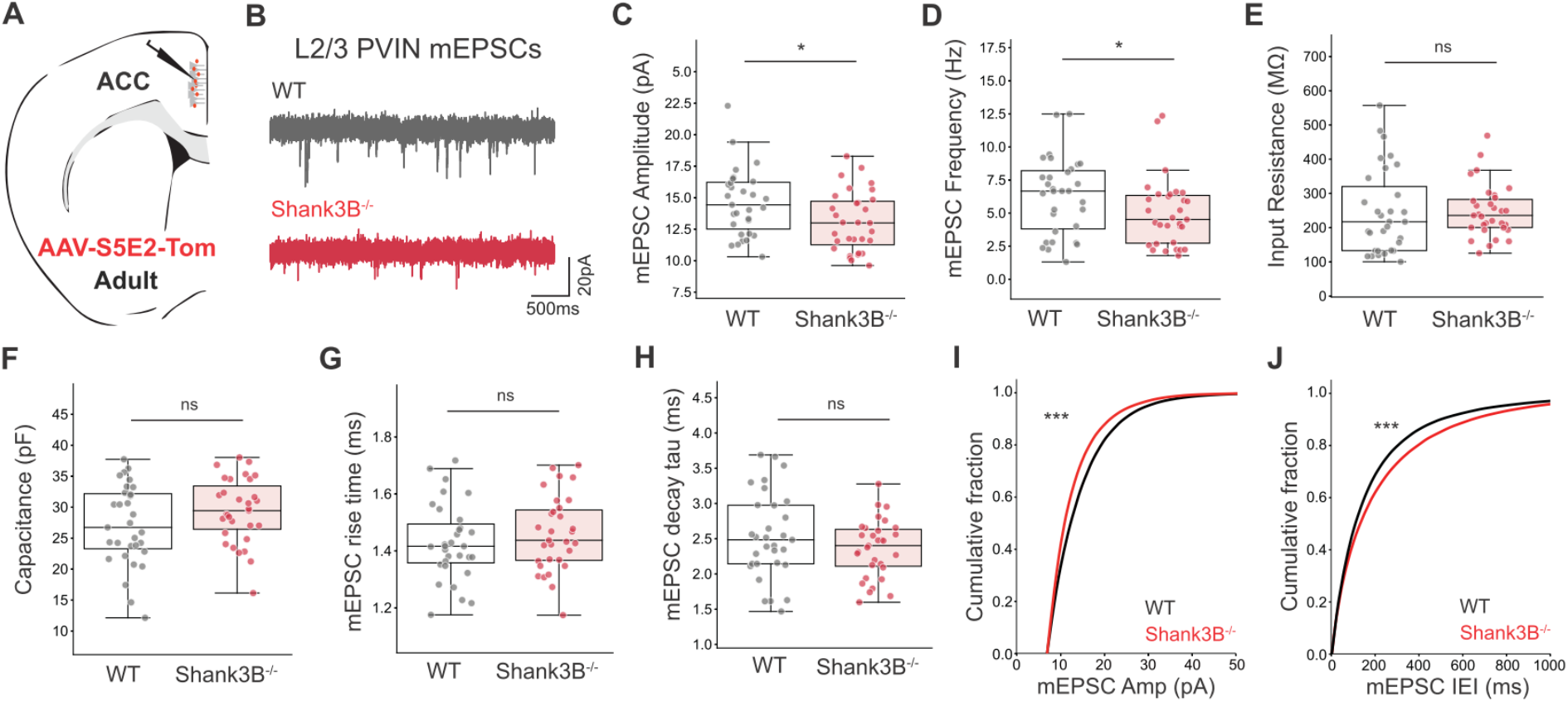
Reduced mEPSC frequency and amplitude in PVINs of L2/3 ACC in adult Shank3B^−/−^ mice. (**A**) Schematic representing coronal brain section and whole-cell recordings in L2/3 ACC on adult mice previously injected with AAV9-S5E2-tdTomato to label PVINs (**B**) Representative AMPAR mEPSCs in L2/3 ACC PVINs of adult WT and Shank3B^−/−^ mice. (**C**) Average ± SD of mEPSC amplitude and (**D**) frequency (**E**) input resistance **(F)** membrane capacitance and **(G-H)** mEPSC rise and decay times of WT (gray) and Shank3B^−/−^ (red) neurons. (**I-J**) Cumulative distribution of mEPSC amplitude. and inter-event interval. n = 31,31 neurons from n = 5,4 WT and Shank3B^−/−^ mice. Box plots show median values (middle vertical bar) and 25th (bottom line of the box) and 75th percentiles (top line of the box) with whiskers indicating the range. eCDFs represent the average of distributions of 100 randomly sampled mEPSCs from each recorded neuron. Average mEPSC box plot *P < 0.05 with Mann–Whitney U test. mEPSC eCDFs ***P < 0.001 with Kolmogorov-Smirnov test.

### Late onset of mEPSC deficits in PYR and PVINs of L2/3 ACC *Shank3B^−/−^* mice

*Shank3B^−/−^* mice already exhibit cortical hyperactivity during early postnatal development (Peixoto et al. 2016) and many of the behavioral deficits in these animals are already observed at P15-P21 (Peixoto et al. 2019), suggesting that neural phenotypes observed at this age are important pathogenic mechanisms with significant therapeutic potential. In addition, glutamatergic innervation of striatal projection neurons in *Shank3B^−/−^*mice shows a biphasic developmental trajectory characterized by early increase and subsequent reduction of mEPSC frequency (Peça et al. 2011; Peixoto et al. 2016), revealing potential developmental compensations in connectivity or function of cortical circuits upon loss of Shank3. To determine whether the mEPSC abnormalities observed in L2/3 ACC PYR and PVIN of adult *Shank3B^−/−^* mice are already observed during postnatal periods, we recorded mEPSCs from the same populations at P15 (Figure 4). Notably, and contrary to what is observed in adult stages, we observed no difference in the amplitude of AMPAR mEPSCs in PYR neurons at P15 (Figure 4C and 4I, Average ± SD PYR mEPSC amplitude at P15, WT 10.6 ± 1.0 pA; KO 10.7 ± 1.3 pA, unpaired Mann Whitney test p = 0.955. Kolmogorov-Smirnov test p = 0.072, n = 100 mEPSCs per n= 42,35 neurons from n = 4,3 WT and Shank3B^−/−^ mice). In addition, Shank3B^−/−^ PYR neurons showed no difference in average mEPSC frequency compared to WT, but a statistically significant increase of individual mEPSC IEI following eCDF distribution analysis (Figure 4D and 4J, Average PYR mEPSC frequency at P15, WT 2.9 ± 1.8 Hz; KO 3.1 ± 1.7 Hz, unpaired Mann Whitney test p = 0.44. Kolmogorov-Smirnov test p = 0.007), suggesting a modest increase in number of glutamatergic synapses. No differences in mEPSC rise and decay times, membrane capacitance, and membrane resistance were detected in P15 Shank3B^−/−^ PYR neurons compared to WT controls (Figure 4E - 4H, average PYR input resistance, WT 186.2 ± 39.8 MΩ; KO 178.8 ± 43.2 MΩ, unpaired Mann Whitney test p = 0.399. Average PYR membrane capacitance, WT 59.1 ± 12.9 pF; KO 57.9 ± 11.4 pF, unpaired Mann Whitney test p = 0.771. Average PYR mEPSC rise time, WT 2.1 ± 0.1 ms; KO 2.1 ± 0.2 ms, unpaired Mann Whitney test p = 0.309. Average PYR mEPSC decay tau, WT 6.0 ± 0.6 ms; KO 5.7 ± 0.7 ms, unpaired Mann Whitney test p = 0.131). These findings indicate that the lower mEPSC frequency and amplitude observed in mature Shank3B^−/−^ PYR neurons only emerge later in development.

**Fig 4.**
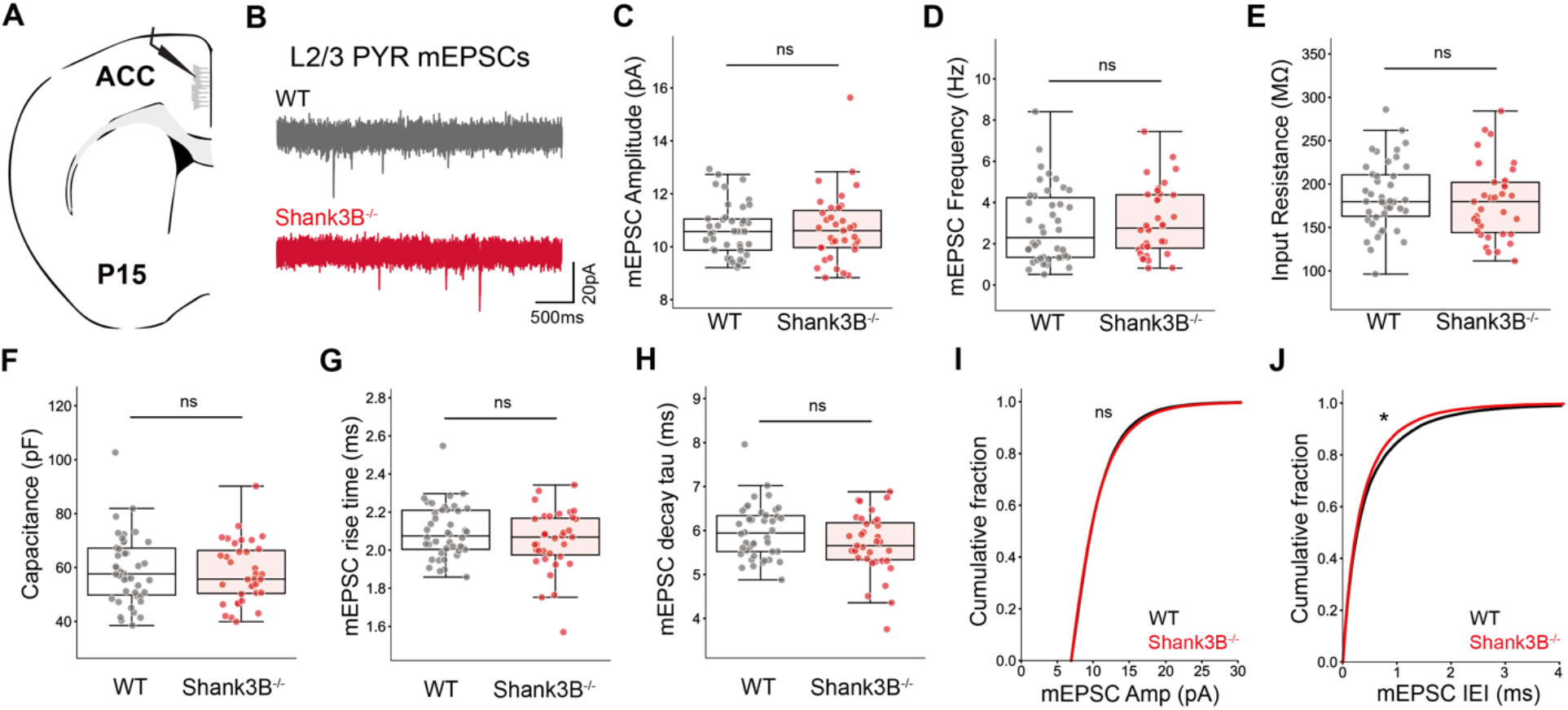
Normal AMPAR mEPSCs of L2/3 PYR in ACC of P15 Shank3B^−/−^ mice. (**A**) Schematic representing coronal brain section and whole-cell recordings in L2/3 ACC of P15 mice (**B**) Representative AMPAR mEPSCs in L2/3 ACC PYR neurons of WT and Shank3B^−/−^ mice. (**C**) Average ± SD of mEPSC amplitude and (**D**) frequency (**E**) input resistance **(F)** membrane capacitance and **(G-H)** mEPSC rise and decay times of WT (gray) and Shank3B^−/−^ (red) neurons. (**I-J**) Cumulative distribution of mEPSC amplitude and inter-event interval. n = 42,35 neurons from n = 4,3 WT and Shank3B^−/−^ mice. Box plots show median values (middle vertical bar) and 25th (bottom line of the box) and 75th percentiles (top line of the box) with whiskers indicating the range. eCDFs represent the average of distributions of 100 randomly sampled mEPSCs from each recorded neuron. eCDF *P < 0.05 with Kolmogorov-Smirnov test.

To determine whether early deficits in glutamatergic innervation of PVINs contribute to the cortical hyperactivity observed in postnatal *Shank3B^−/−^* mice, we characterized mEPSC in L2/3 PVINs in P15 ACC after infection with AAV9-S5E2-tdTom at P7 (Figure 5). This strategy results in labeling of 82.4% PV neurons in ACC at P21 (Figure 2B and 2D, Average ± SEM Fraction of S5E2-tdTom^+^ neurons with PV^+^ signal 82.4 ± 4.1%, n = 132 cells from 2 WT), consistent with previous reports (Vormstein-Schneider et al. 2020). Similar to what we observed in L2/3 PYR neurons, there was no statistically significant difference of mEPSC amplitude and frequency between WT and Shank3B^−/−^ L2/3 ACC PVINs during this early postnatal period (Figure 5C and 5D, Average ± SD PVIN mEPSC amplitude at P15, WT 20.1 ± 4.1 pA; KO 19.8 ± 4.3 pA, unpaired Mann Whitney test p = 0.536. Average PVIN mEPSC frequency in adult, WT 7.8 ± 3.0 Hz; KO 8.4 ± 3.5 Hz, unpaired Mann Whitney test p = 0.348. Figure 5I and 5J, Kolmogorov-Smirnov test p = 0.48 and 0.32, n = 100 mEPSCs per n= 69,65 neurons from n = 6,6 WT and Shank3B^−/−^ mice). Likewise, WT and Shank3B^−/−^ PVINs showed comparable mEPSC rise and decay times, and membrane capacitance and resistance (Figure 5E-5H, Average ± SD PVIN input resistance, WT 247.8 ± 98.9 MΩ; KO 259.5 ± 108.6 MΩ, unpaired Mann Whitney test p = 0.413. Average ± SD PVIN membrane capacitance, WT 41.1 ± 8.8 pF; KO 41.2 ± 7.6 pF, unpaired Mann Whitney test p = 0.936. PVIN mEPSC rise time, WT 1.5 ± 0.1 ms; KO 1.5 ± 0.1 ms, unpaired Mann Whitney test p = 0.468. PVIN mEPSC decay tau, WT 3.2 ± 0.5 ms; KO 3.2 ± 0.4 ms, unpaired Mann Whitney test p = 0.918). Together, these results indicate that AMPAR inputs onto both L2/3 ACC PYR and PVINs in Shank3B^−/−^ mice are not yet perturbed at P15, and that deficits in mEPSC observed later in adulthood arise after the ∼P15 postnatal period, potentially as compensatory mechanisms (Tatavarty et al. 2020).

**Fig 5.**
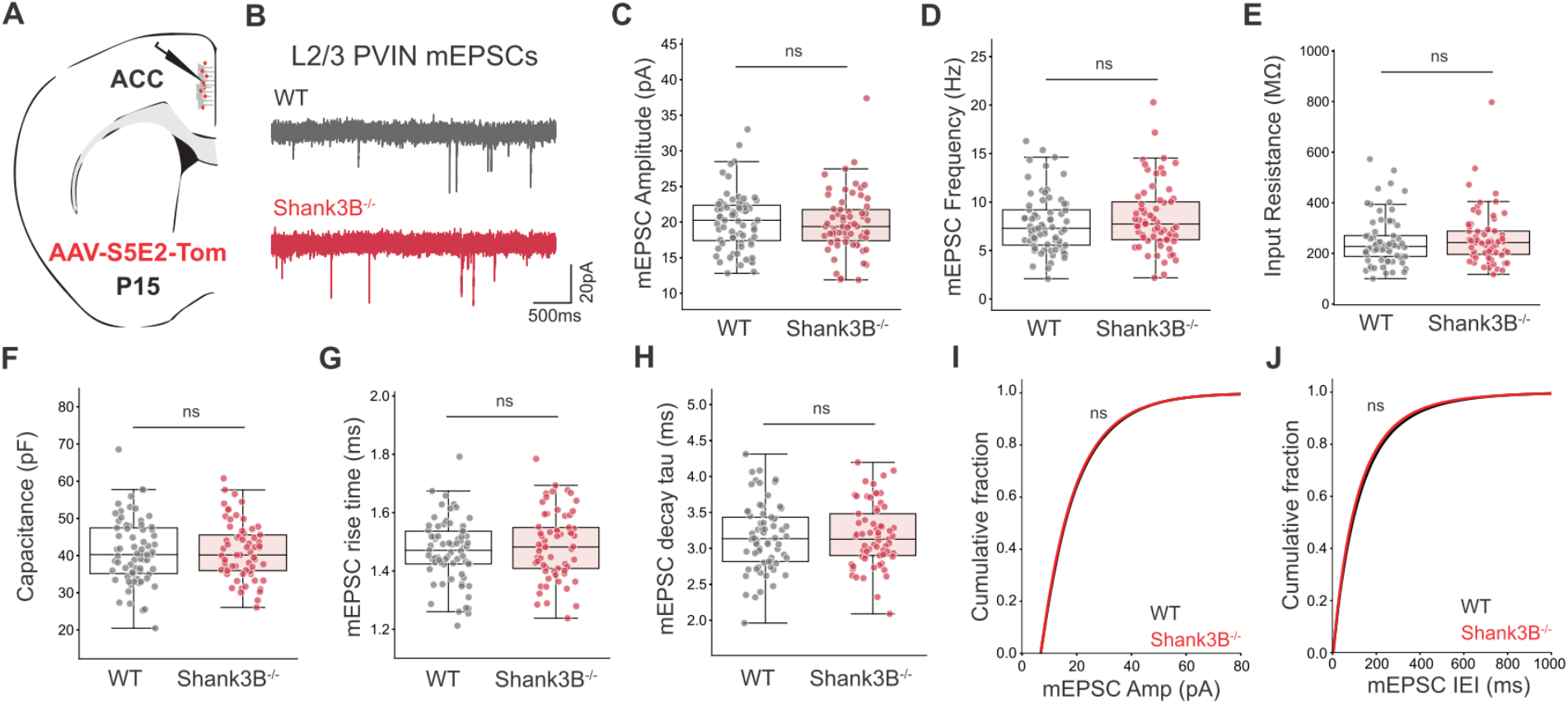
Normal AMPAR mEPSCs of L2/3 PVINs in ACC of P15 Shank3B^−/−^ mice. (**A**) Schematic representing coronal brain section and whole-cell recordings in L2/3 ACC on adult mice previously injected with AAV8-S5E2-tdTomato to label PVINs (**B**) Representative AMPAR mEPSCs in L2/3 ACC PVINs of P15 WT and Shank3B^−/−^ mice. (**C**) Average ± SD of mEPSC amplitude and (**D**) frequency (**E**) input resistance **(F)** membrane capacitance and **(G-H)** mEPSC rise and decay times of WT (gray) and Shank3B^−/−^ (red) neurons. (**I-J**) Cumulative distribution of mEPSC amplitude and inter-event interval. n = 69,65 neurons from n = 6,6 WT and Shank3B^−/−^ mice. Box plots show median values (middle vertical bar) and 25th (bottom line of the box) and 75th percentiles (top line of the box) with whiskers indicating the range. eCDFs represent the average of distributions of 100 randomly sampled mEPSCs from each recorded neuron.

### Regional heterogeneity of mEPSC deficits in L2/3 cortical PVINs in adult *Shank3B^−/−^* mice

Loss of Shank3 leads to hypoactivity of Dlx5/6-positive GABAergic interneurons in L2/3 of adult primary somatosensory (S1) cortex (Chen et al. 2020) and *Shank3B^−/−^* mice exhibit abnormal expression patterns of PV across multiple cortical regions (Deemyad et al. 2021; Filice et al. 2016, 2020; Gogolla et al. 2014; Orefice et al. 2019). Given that PVINs of L2/3 ACC show reduced mEPSC frequency and amplitude in adulthood (Figure 3) we determined whether PVINs of L2/3 S1 also show deficits in AMPAR mEPSCs, as that could underlie the abnormal responses to sensory stimuli observed in S1 cortex of *Shank3B^−/−^* mice (Figure 7). Infection with AAV9-S5E2-tdTom at P45 results in high specificity of PVIN labeling in P60 S1 somatosensory cortex (Figure 6, Average ± SEM Fraction of S5E2-tdTom^+^ neurons with PV^+^ co-localization 92.3 ± 3.1%, n = 181 cells from 3 WT).

**Fig 6.**
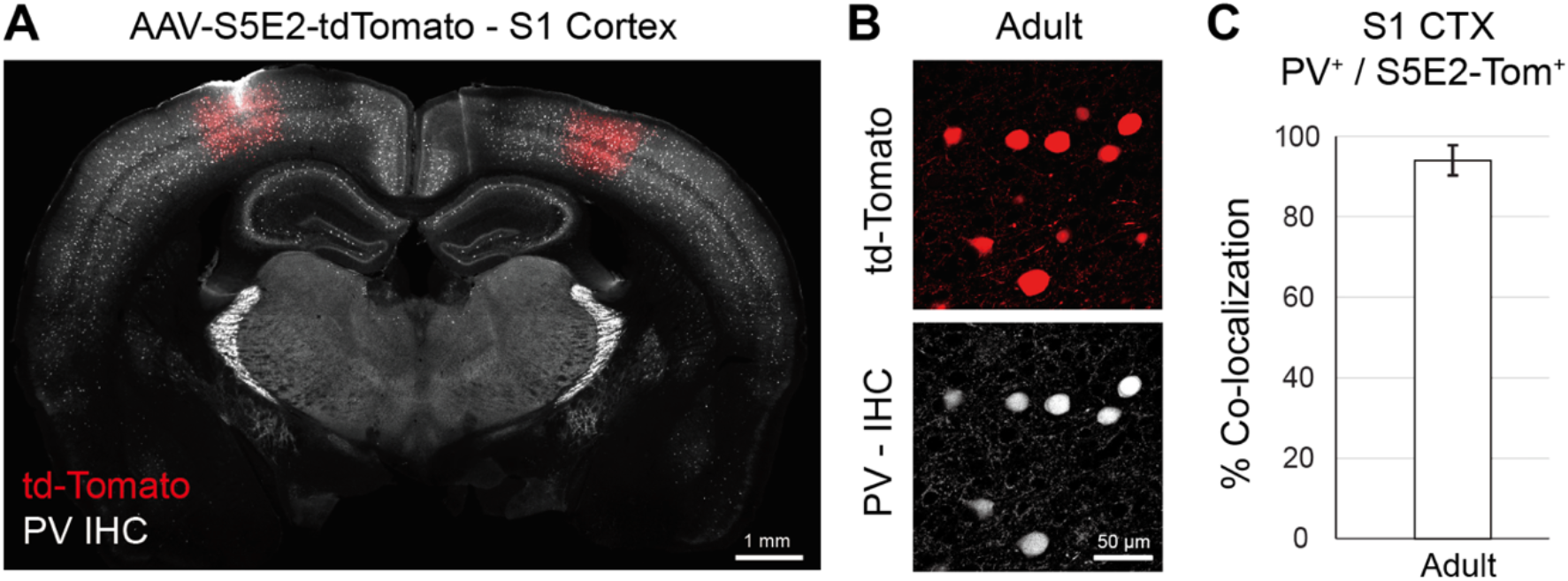
Specific labeling of PVINs with AAV9-S5E2-tdTom in S1 cortex of adult mice. (**A**) Representative image of PV IHC in S1 of P60 WT mouse injected with AAV9-S5E2-tdTom at P45 (**B**) Magnified images of PV IHC after 2 weeks of injection (**C**) Percentage of tdTom-positive cells with PV co-labeling - Adult S1 n = 181 cells from n = 2 mice.

**Fig 7.**
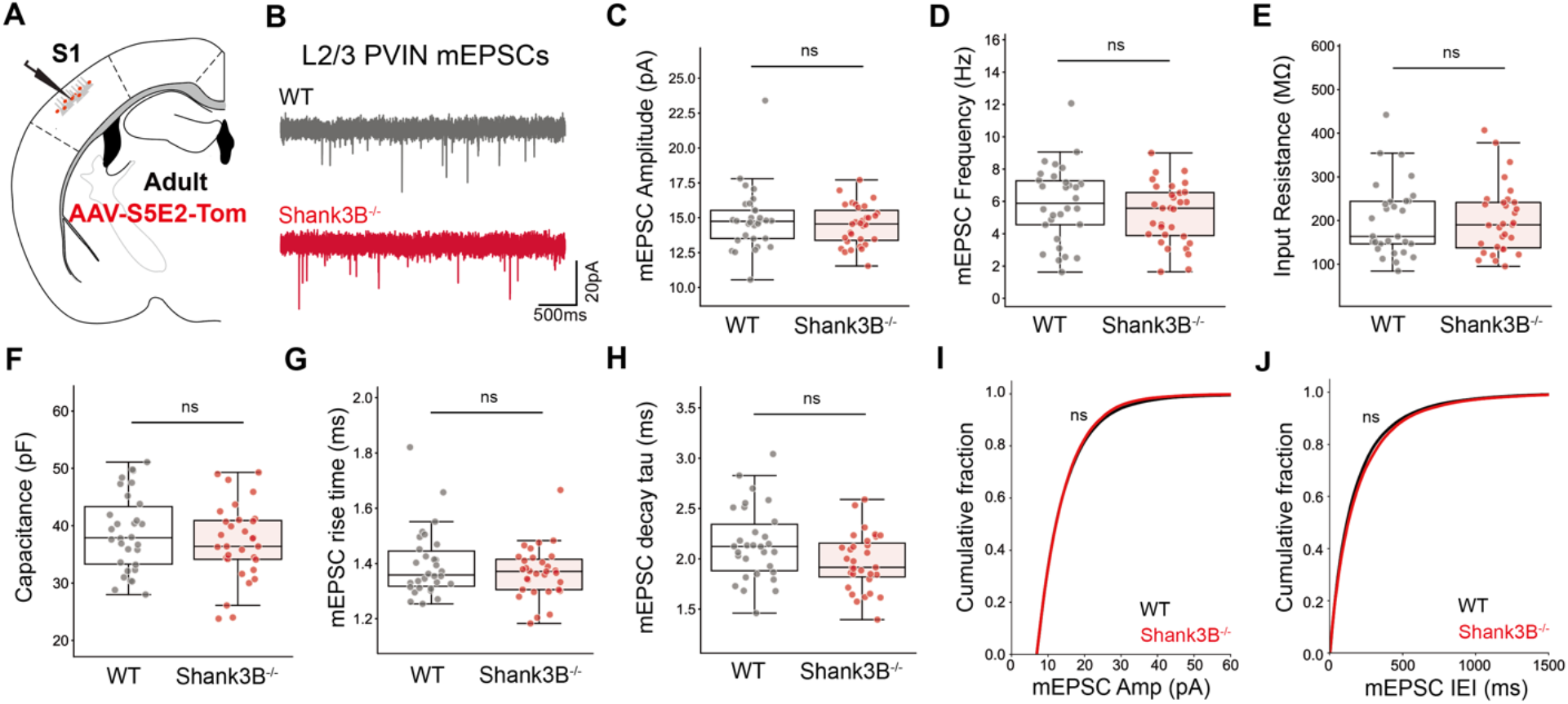
No excitatory synaptic dysfunction of L2/3 PVIN in S1 is found in adult Shank3B^−/−^ animals. (**A**) Schematic showing mEPSC recording in L2/3 PVIN in adult S1. (**B**) Representative AMPAR mEPSC recordings in L2/3 PVIN in S1 of WT and Shank3B^−/−^ mice in adult. (**C**) No statistic difference of mEPSC amplitude was found in L2/3 PVIN in adult Shank3B^−/−^ S1. (**D**) No statistic difference of mEPSC frequency of L2/3 PVIN in adult S1 was found. (**E, F**) Shank3B^−/−^ showed no effect on both membrane capacitance and input resistance of L2/3 PVIN in adult S1. (**G, H**) No statistic difference was found in mEPSC rise time and decay tau. (**I**) Cumulative distribution of mEPSC amplitude. (**J**) Cumulative distribution of mEPSC inter-event interval. n = 30 neurons from 3 WT mice; 31 neurons from 3 Shank3B^−/−^ mice. Box plots show median values (middle vertical bar) and 25th (bottom line of the box) and 75th percentiles (top line of the box) with whiskers indicating the range. eCDFs represent the average of distributions of 100 randomly sampled mEPSCs from each recorded neuron. *P < 0.05 with Mann–Whitney U test. *P < 0.05 with Kolmogorov-Smirnov test.

Surprisingly, we found no difference in mEPSC amplitude and frequency between adult Shank3B^−/−^ and WT PVINs of L2/3 S1 cortex (Figure 7C and 7D, average PVIN mEPSC amplitude in adult, WT 14.8 ± 2.3 pA; KO 14.5 ± 1.5 pA, unpaired Mann Whitney test p = 0.639. Average ± SD PVIN mEPSC frequency, WT 5.9 ± 2.4 Hz; KO 5.2 ± 1.9 Hz, unpaired Mann Whitney test p = 0.296. Figure 7I and 7J, Kolmogorov-Smirnov test p = 0.96 and 0.18, n = 100 mEPSCs per n= 30,31 neurons from n = 3,3 WT and Shank3B^−/−^ mice). Moreover, Shank3B^−/−^ PVINs exhibited normal AMPAR mEPSC kinetics, as well as membrane capacitance and input resistance (Figure 7D - 7H, Average ± SD PVIN input resistance, WT 201.2 ± 84.4 MΩ; KO 202.0 ± 78.3 MΩ, unpaired Mann Whitney test p = 0.9.7; Membrane capacitance, WT 38.8 ± 6.7 pF; KO 37.1 ± 6.5 pF, unpaired Mann Whitney test p = 0.462; mEPSC rise time, WT 1.4 ± 0.1 ms; KO 1.4 ± 0.1 ms, unpaired Mann Whitney test p = 0.639; mEPSC decay tau, WT 2.1 ± 0.4 ms; KO 2.0 ± 0.3 ms, unpaired Mann Whitney test p = 0.064). Together, these findings indicate that L2/3 PVINs are differentially affected across sensory and frontal cortical regions and suggest that AMPAR synaptic dysfunction is not a critical pathophysiology underlying the abnormal tactile hyperexcitability observed in the adult barrel cortex of *Shank3B^−/−^* mice.

## Discussion

Despite the growing prevalence of ASDs and their enormous disease burden, no effective pharmacological treatments have been developed for these conditions (Jeste and Geschwind 2016). Failure to identify successful therapeutic targets has been attributed to the high genetic heterogeneity and variable clinical presentation across the autistic spectrum (Lord et al. 2000; De Rubeis et al. 2018). However, the distinctive developmental trajectory and selective deterioration of specific motor and cognitive abilities presented by autistic children suggest the potential implication of common pathogenic processes that disrupt convergent developmental processes during critical periods of brain development (Klei et al. 2012; Pinto et al. 2014). Identification of such mechanisms would represent a long-needed breakthrough for the development of new therapies.

Here, we characterized the maturation of glutamatergic connectivity in PYR and PVINs of ACC, a region of mPFC that integrates information from limbic, sensory, and motor circuits to mediate sensory processing, social cognition and goal directed behaviors, functions critically impaired in ASDs (Apps, Rushworth, and Chang 2016; Balsters et al. 2017; Stevens, Hurley, and Taber 2011; Zhou et al. 2016). PVINs regulate local cortical circuits by mediating feedforward inhibition and gamma oscillations and their activity depends on precise spatiotemporal integration of excitatory inputs (Akgül and McBain 2016; Jonas et al. 2004). Shank3 regulates the postnatal maturation of excitatory synapses by controlling the trafficking and retention of AMPAR subunits either directly or through association with other scaffolding proteins (Bariselli et al. 2016; Chiola et al. 2021; Mameza et al. 2013; Raynaud et al. 2013; Verpelli et al. 2011). We thus hypothesized that depressed glutamatergic connectivity in PVINs could be a potential pathophysiological phenotype associated with the early cortical hyperactivity and behavioral deficits observed in postnatal Shank3B^−/−^ mice (Peixoto et al. 2016, 2019). However, we detected no differences in AMPAR mEPSC frequency and amplitude of both L2/3 PYR and PVINs at P15 (Figures 4 and 5), suggesting that glutamatergic connectivity in these two neural populations is not affected by loss of Shank3 during this early developmental stage. This in an interesting and important finding because it shows that glutamatergic synapse deficits in these neurons are not a primary pathogenic factor in Shank3B^−/−^ mice. By contrast, ACC PYR neurons of adult Shank3B^−/−^ mice exhibited reduced AMPAR mEPSC amplitude, consistent with depressed AMPAR synaptic function. Notably, we did not observe a reduction in mEPSC frequency in adult ACC PYR neurons as reported in a previous study (Guo et al. 2019). One possible explanation for this discrepancy is that we employed more stringent mEPSC detection methods that exclude low amplitude events, which might have accounted for the high frequencies (∼12Hz) and low amplitude (∼8pA) mEPSCs reported in that study. In fact, eCDF analysis of individual mEPSCs revealed a statistically significant increase in mEPSC frequency that is consistent with another study that characterized L2/3 PYR neurons in mPFC of a different SHANK3 deletion (exon 14-16) mouse line (Yoo et al. 2019). Moreover, membrane resistance of PYR neurons was increased, suggesting a complex pathophysiology of L2/3 PYR neurons characterized by depressed synaptic function but increased excitability. Future work characterizing the *in vivo* functional implication of these phenotypes will be important to understand their pathophysiological role. Nevertheless, our findings support that loss of Shank3 leads to abnormal synaptic function of ACC PYR neurons, but these abnormal phenotypes only emerge later in development.

In perhaps the most important contribution of this work, we found robust downregulation of both mEPSC frequency and amplitude in L2/3 ACC PVINs in adulthood, providing the first demonstration of synaptic deficits in PVINs of Shank3B^−/−^ mice. Notably, Shank3B^−/−^ and numerous other ASD mouse models show reduced of PV expression in cortex (Deemyad et al. 2021; Filice et al. 2016; Gogolla et al. 2014; Negwer et al. 2020; Orefice et al. 2016, 2019; Peñagarikano et al. 2011; Pinto et al. 2014; Vogt et al. 2018; Wen et al. 2019; Zhang et al. 2020). Enhancing GABA function or inhibiting glutamatergic activity improves abnormal behaviors in some of these models, suggesting that interventions normalizing cortical activity imbalances hold potential therapeutic value (Han et al. 2012, 2014; Kim et al. 2017). However, developmental expression of PV itself is regulated by network activity (Alcántara et al. 1996; Patz et al. 2004) which raises the question of whether abnormal PV levels associated with ASD represent a primary pathophysiology or a secondary developmental compensation. Our results showing normal AMPAR synaptic connectivity in PVINs early in development (Figure 5) suggest the latter, but do not exclude the possibility that loss of Shank3 affects other physiological properties not revealed by our mEPSC recordings. Interestingly, we did not observe mEPSC deficits in PVINs of the somatosensory cortex of adult Shank3B^−/−^ mice (Figure 6), which was surprising because Dlx5/6-positive interneurons in the barrel cortex of Shank3B^−/−^ mice show reduced activity in response to whisker simulation (Chen et al. 2020). Again, our data suggests these deficits are not caused by reduced glutamatergic inputs in PVINs but does not exclude the potential presence of other physiological abnormalities (Orefice et al. 2016). Another possibility is that lower activity observed in the Dlx5/6 neuronal population of S1 cortex is mediated by deficits in Somatostatin interneurons, as they are also labeled by Dlx5/6 promoter-based tools and are affected by SHANK3 mutations (Ali et al. 2021). Regardless, our data indicates that PVINs of different cortical regions are differently affected by loss of Shank3. This regional heterogeneity is consistent with the variable pattern of PV expression reduction across the cortex of Shank3B^−/−^ mice (Deemyad et al. 2021) and suggests that some cortical regions might be more susceptible to these conditions.

Most individuals with *SHANK3* disorders develop seizures during their lifetimes (Kolevzon et al. 2019; De Rubeis et al. 2018) and many exhibit patterns of epileptiform activity in EEG recordings before seizure onset (Khan et al. 2018). Anticonvulsants and benzodiazepines are often administered after seizure onset but are not commonly used prophylactically (de Coo et al. 2023; Guilmatre et al. 2014). Further understanding the mechanisms underlying cortical hyperactivity and PVIN dysfunction associated with SHANK3 disorders will provide valuable information to design optimal early interventions that correct, and not exacerbate, cortical circuit defects associated with these conditions.

## Acknowledgements

We thank Tasha Merchant, Lan Chen, Sidney Dawkins and Andrew D’Agostino for assistance with mouse husbandry and genotyping. R.T.P was supported by R01MH124695, R21MH132015 and a Bridge to Independent Award from the Simons Foundation Autism Research Initiative. L.N. was supported by T32 MH16804.

## Methods

### Animals

All experimental manipulations on mice were performed in accordance with protocols approved by the Institutional Animal Use and Care Committee at the University of Pittsburgh in compliance with the guidelines described in the US National Institutes of Health *Guide for the Care and Use of Laboratory Animals*. Mice were housed on a 12/12hr light/dark cycle with chow and water provided ad libitum. Mice were weaned at P21-23 and separated by sex in cages of 2-5 animals of mixed genotypes. Shank3B^−/−^ mutant mice were described previously (Peça et al. 2011) and obtained from The Jackson Laboratory (#017688). Characterization of neural properties by electrophysiology was performed in Shank3B^+/+^ and Shank3B^−/−^ age-matched mice from breeding pairs between Shank3B^+/−^ heterozygous animals. Male mice were used in all experiments as no sexual dimorphism was ever detected in the mouse lines used and behavioral assays tested or reported in previous studies.

### Viruses and stereotaxic intracranial injections

Intracranial injections were performed at age P7-P8 or 2 weeks before recording for P15 and adult experiments, respectively. Mice were anesthetized with isoflurane (4% for induction and 1-2% thereafter for maintenance) placed into a stereotaxic apparatus (Kopf Instruments model 1900). Mice received carprofen for analgesia before surgery. The hair on head of adult mice was removed by hair removal creams (Nair). The surgical area was cleaned by 2% iodine and 70% alcohol solution. To perform intracranial injections, injection sites were poked by sterile 30G needle in pups and drilled by Micromotor drill (Stoelting) in adults, respectively. All skull measurements were made relative to Bregma, and viruses were delivered by injecting 350 nl at a maximum rate of 75 nL/min using a Harvard Apparatus PHD ULTRA CP injector. For labeling of PVINs, virus expressing tdTomato under the PVIN specific S5E2 enhancer (AAV9-S5E2-tdTomato, Addgene 135630-AAV9) was injected in mice using coordinates: AP +0.5 mm; ML 0.3 mm; Depth −1.0 mm for pups and AP +1.1 mm; ML 0.35 mm; Depth −1.1 mm for adults. Following injections and wound suture, mice were placed on heating pad till awakening and returned to home cages for at least 7 days.

### Brain slice preparation and whole-cell electrophysiology

Acute brain slices were prepared following anesthesia by isoflurane inhalation and transcardiac perfusion with ice-cold artificial cerebrospinal fluid (ACSF) containing (in mM): 125 NaCl, 2.5 KCl, 25 NaHCO_3_, 2 CaCl_2_, 1 MgCl_2_, 1.25 NaH_2_PO_4_ and 25 glucose (310 mOsm per kg). Cerebral hemispheres were removed and transferred into a slicing chamber containing ice-cold ACSF. Coronal slices including ACC or S1 (275 µm thick) were cut with a Leica VT1200s vibratome and transferred for 10 min to a holding chamber containing choline-based solution consisting of (in mM): 110 choline chloride, 25 NaHCO_3_, 2.5 KCl, 7 MgCl_2_, 0.5 CaCl_2_, 1.25 NaH_2_PO_4_, 25 glucose, 11.6 ascorbic acid, and 3.1 pyruvic acid at 33°C. Slices were subsequently transferred to a chamber with pre-warmed ACSF and maintained at room temperature (20–22°C). All recordings were obtained within 4 hours of slicing. Both ACSF and choline solution were constantly bubbled with 95% O_2_/5% CO_2_. Individual slices were transferred to a recording chamber mounted on an upright microscope (Olympus BX51WI) and continuously perfused (1–2 ml per minute) with ACSF at room temperature. Cells were visualized using a 40× water-immersion objective with infrared illumination. Whole-cell voltage clamp recordings were made from pyramidal neurons or PVINs of ACC or S1 cortex. Patch pipettes (2–4 MΩ) pulled from borosilicate glass (BF150-86-7.5, Sutter Instruments) were filled with a Cs^+^-based internal solution containing (in mM): 130 CsMeSO_4_, 10 HEPES, 1.8 MgCl_2_, 4 Na_2_ATP, 0.3 NaGTP, and 8 Na_2_-phosphocreatine,10 CsCl_2,_ 3.3 QX-314 (Cl^−^ salt), (pH 7.3 adjusted with CsOH; 300 mOsm per kg) for voltage-clamp recordings. For all voltage-clamp experiments, errors due to voltage drop across the series resistance (<20 MΩ) were left uncompensated. For mEPSC recordings, ACSF contained 1 μM TTX, 1 μM (RS)-CPP, and 1 μM Gabazine, and recordings were performed at room temperature (20-22°C) with Vm = −70mV. After breaking in, cells were left to stabilize for 4 min and currents were then acquired continuously for 5 min. Membrane currents and potentials were amplified and low-pass filtered at 3 kHz using Multiclamp 700B amplifier (Molecular Devices), digitized at 10 kHz, and acquired using National Instruments acquisition boards and a custom version of ScanImage written in MATLAB (Mathworks). Calculation of input resistance and membrane capacitance were performed by fitting evoked currents in response to −5mV voltage steps in the first seconds after cell break-in. Data saved as Matlab files for subsequent off-line analysis performed by using custom Python program ClampSuite. The software package is available at github.com/LarsHenrikNelson/ClampSuite.

### Brain tissue imaging and immunohistochemistry

Mice were deeply anesthetized with isofluorane and perfused transcardially with 1X phosphate-buffered saline (PBS) (pH 7.4) followed by 4% paraformaldehyde in PBS. Brains were fixed overnight at 4°C, washed in phosphate buffer saline (PBS) and sectioned (50 µm) coronally using a vibratome (Leica VT1000s). Brain sections were incubated in blocking buffer containing: 10% normal goat serum, mouse IgG (1 μg/mL) and 0.3% triton X-100 in 1X PBS for one hour at room temperature. After blocking, sections were incubated overnight with anti-Parvalbumin antibodies (1:10000, SWant PV235) at 4°C. After the sections were washed, they were incubated with anti-mouse Alexa 647 (1:2000, Abcam) and DAPI (1 μg/mL, Thermo Fisher) for 1-2 hours at room temperature. The sections were mounted on glass slides, dried and mounted with ProLong antifade reagent containing DAPI (Molecular Probes). Whole brain sections were imaged by an Olympus FV1200 confocal microscope with 60X objective lens. For quantification of PV-tdTom co-localization three slices containing relevant cortical regions were analyzed per animal to quantify percentage of PV-positive cells among S5E2-tdTom labeled population. For each slice three regions were randomly selected and imaged and cell number was manually quantified. PV signal was considered positive if fluorescence was higher that 2.5x background intensity. A population of S5E2-tdTom-labeled cells with very small cell size (<100 μm^2^) in adult ACC was excluded from quantification and whole-cell patch clamp recordings.

### Data and statistical analyses

No statistical methods were used to predetermine sample sizes, but our sample sizes are similar to those reported in previous in previous publications (Kozorovitskiy et al. 2015; Peixoto et al. 2012, 2016, 2019). With few exceptions, hypothesis testing between two groups use Mann– Whitney U test that does not assume the underlying data are normally distributed. Kolmogorov-Smirnov test was used for comparing cumulative distribution between two groups (add details about sampling). Values are expressed as the median values (middle vertical bar) and 25^th^ (bottom line of the box) and 75th percentiles (top line of the box) with whiskers indicating the range in box plot. Statistical significance was accepted when P < 0.05. Calculation of the statistics was done in Scipy (Python). Details of particular statistical analyses can be found in the respective figures/results section for each dataset. mEPSCc and mIPSCs were analyzed using a custom python-based program Clampsuite available at https://github.com/LarsHenrikNelson/ClampSuite. Acquisition offset was removed by subtracting the mean. Recordings were filtered using a zero-phase Remez filter with a low pass filter at (PYR - 600/300Hz; PVIN - 600/600Hz). Events were identified using a modified FFT deconvolution method described in (Pernía-Andrade et al. 2012). Events were excluded based on the following criteria: PYR - Amplitude lower than 7 pA, rise time > 0.9ms and <4ms, minimum decay time of 4 ms and a minimum interevent distance of 2 ms; PVIN - Amplitude lower than 7 pA, rise time > 0.8ms and < 4ms, minimum decay time of 0.6 ms and a minimum interevent distance of 2 ms; Tau was estimated using the equation (event amplitude) * (1 / e1)) and the time when the event reached tau was interpolated from the time array associated with the event decay.

